# Hydrogen-deuterium exchange mass spectrometry captures distinct dynamics upon substrate and inhibitor binding to a transporter

**DOI:** 10.1101/2020.04.03.023564

**Authors:** Ruyu Jia, Chloe Martens, Mrinal Shekhar, Shashank Pant, Grant A. Pellowe, Andy M. Lau, Heather E. Findlay, Nicola J. Harris, Emad Tajkhorshid, Paula J. Booth, Argyris Politis

## Abstract

Proton-coupled transporters use transmembrane proton gradients to power active transport of nutrients inside the cell. High-resolution structures often fail to capture the coupling between proton and ligand binding, and conformational changes associated with transport. We combine HDX-MS with mutagenesis and MD simulations to dissect the molecular mechanism of the prototypical transporter XylE. We show that protonation of a conserved aspartate triggers conformational transition from outward-facing to inward-facing state. This transition only occurs in the presence of substrate xylose, while the inhibitor glucose locks the transporter in the outward-facing state. MD simulations corroborate the experiments by showing that only the combination of protonation and xylose binding, and not glucose, sets up the transporter for conformational switch. Overall, we demonstrate the unique ability of HDX-MS to distinguish between the conformational dynamics of inhibitor and substrate binding, and show that a specific allosteric coupling between substrate binding and protonation is a key step to initiate transport.

## Introduction

Structural biology of membrane proteins has evolved at an increasing pace over the past few years [1]. The more high-resolution structural information becomes available, the clearer it appears that complementary dynamic information is required to understand the mechanism of a protein of interest [2]. Energy coupling in secondary transporters is a good example of the type of information that static structures cannot directly provide about molecular mechanisms [3]. While it is clear that these transporters alternate between different conformations ranging from open to the cytoplasm (inward-facing; IF) to open to the extracellular medium (outward-facing; OF), the molecular chain of events leading to these transitions are difficult to capture [4]. Specifically, the identification of the allosteric networks linking ion and substrate binding, and the ensuing protein conformational changes are difficult to deduce from structural snapshots [5]. Thus, linking structure to mechanism at a molecular level requires characterizing the conformational dynamics of membrane proteins [6].

Among the techniques available to study conformational changes, HDX-MS is a newcomer for the study of membrane proteins [7]. This technique reports on the exchange of amide hydrogens on the protein backbone in the presence of deuterated solvent at a peptide level of resolution [8]. The main advantage over more established methods such as FRET and DEER is that it does not require covalent labelling of the protein of interest, thus bypassing a lot of the molecular biology work and controls [9]. The method also requires lower amount of sample compared to other biophysical methods (such as NMR or X-ray crystallography) [10] and tolerates sample heterogeneity and complexity [11, 12]. H/D exchange however does not strictly report on distance changes involved in conformational transitions. Rather, it reports on the stability of H-bond of the amide backbone which is mainly conditioned by two parameters: local structural dynamics and solvent accessibility [13, 14]. We have shown previously that for a series of transporters, the changes in solvent accessibility can be correlated with conformational changes in most cases [15]. This is particularly helpful to understand the molecular mechanism of transporters as they switch between OF and IF conformations [16]. The conformational effect of ligand binding, mutation of conserved residues or both, can be tested in a systematic way by comparing the H/D exchange pattern in different conditions, in so-called differential ΔHDX-MS experiments. Assuming that no major changes in the stability of transporters occur when introducing either the ligand or a mutation, ΔHDX-MS offers a quick and easy readout of the conformational transition between different states.

The symporter XylE of the ubiquitous MFS family is a bacterial homolog of human glucose transporters and uses the proton-motive force to catalyse xylose translocation across the membrane [17]. The crystal structure of this sugar transporter has been solved in multiple conformations; inward-open, inward-occluded, and outward-occluded with substrate xylose and inhibitor glucose bound [18, 19]. Despite advances, the exact sequence of molecular events enabling the transport cycle and the coupling between xylose and proton binding and conformational changes are not understood. Two transmembrane acidic residues located away from the binding pocket are likely candidates for the initial protonation step: D27 on helix 1 and E206 on helix 6. Biochemical assays have identified D27 as an essential component of active transport and neighbouring E206 has been suggested to play a role in modulating the pK_a_ of D27, to regulate its ability to bind and release proton [20, 21]. Binding assays carried out on wild-type (WT) [18] and D27N mutant [19] show that they both bind xylose with a similar affinity (Kd 0.3 mM). Biochemical studies and MD simulations have shown that protonation must occur before xylose binding but it is not clear whether E206, D27 or both are protonated [19, 22]. Furthermore, the structures of the xylose-bound and glucose-bound protein are virtually identical, with only minor differences in the interaction network at the binding site [23]. This observation raises questions on how the transporter discriminates between substrate and inhibitor and how the potential differences are translated into conformational changes.

In a previous study, we carried out an extensive characterization of the conformational dynamics of XylE by HDX-MS, in order to establish the mechanistic role of a conserved network of charged residues located on the intracellular side [11]. For benchmarking purposes, we locked the transporter in an OF conformation by replacing a conserved glycine necessary for the transition by a bulky tryptophan. This work provided a set of ΔHDX maps associated with transitions toward either the IF or OF states, which can now be used as a fingerprint to guide interpretation of the ΔHDX experiments performed in the present study. Here, we performed HDX-MS measurements of the proton-coupled symporter XylE in the presence of its substrate xylose, inhibitor glucose and mutations at candidate protonation sites D27 and E206 [18, 24]. The systematic HDX analysis coupled to MD simulations identifies differences in structural dynamics and allosteric events between xylose and glucose binding, providing a rationale for inhibitor versus substrate distinction. These findings, in turn, allow us to deduce a likely sequence of events leading to transport.

## Results

To dissect the role of proton and substrate binding, all the possible combinations between WT and mutants mimicking protonation - D27N, E206Q and E206Q&D27N - in the apo and substrate bound states were tested (**Fig. 1A, B)**. At least three biological triplicates were used for each ΔHDX-MS experiment comparing two different protein states [25]. Since the peptides generated by enzymatic digestion can be different between biological replicates, we used Deuteros [26] to identify peptides showing a significant difference in deuterium uptake for each individual ΔHDX-MS experiment and carried out an extra step of curating the data to represent only the peptides that are present in all replicates (**Fig. S4**). It is noted that sequence coverage of more than 90% was obtained in most cases, allowing us to monitor the dynamics of nearly the entire protein.

**Figure 1.**
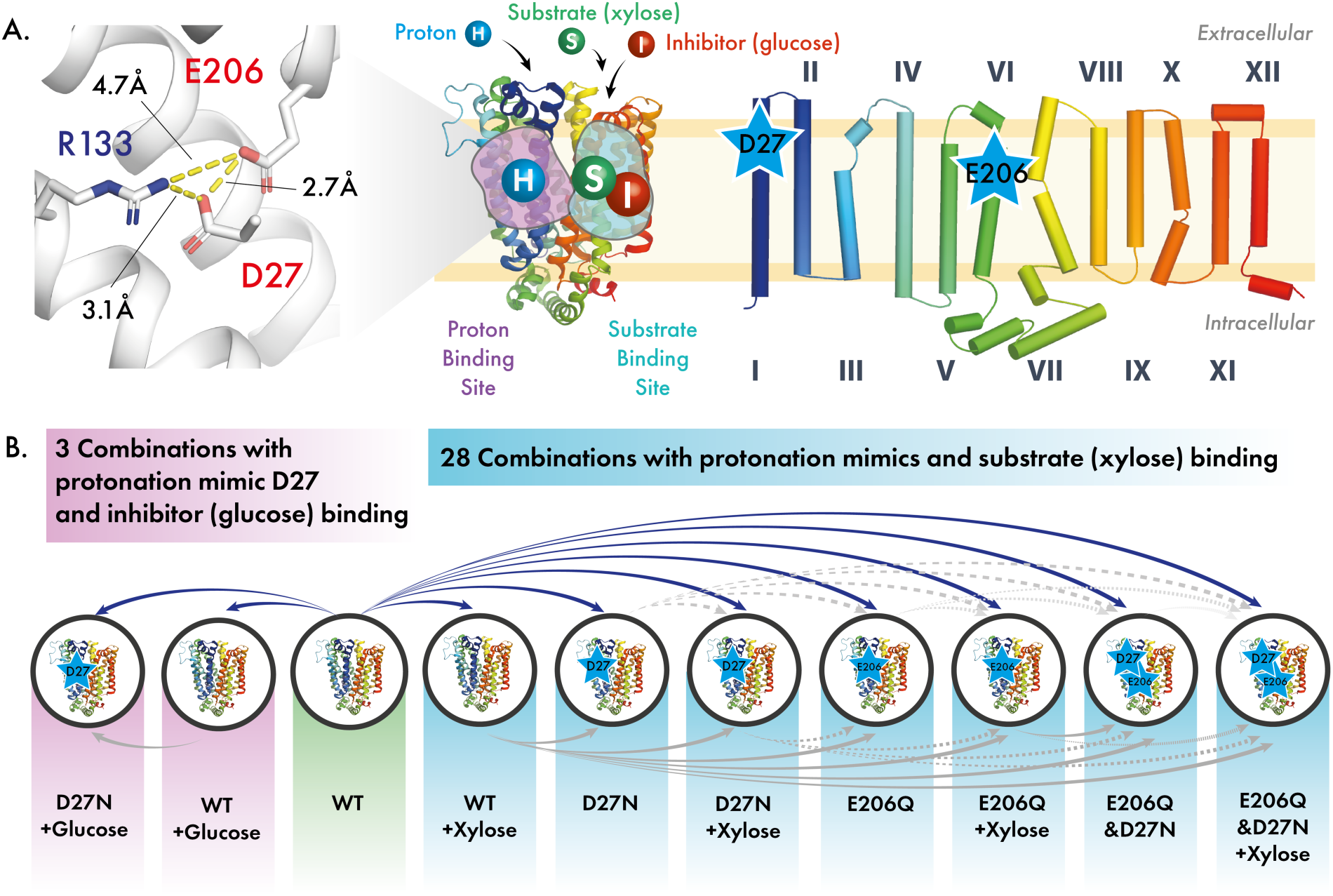
Structure of XylE and proposed experiment design. **(A)**. Topological and 3D structure of XylE (PDB: 4GBY)[18]. Three residues of interest in the proton binding site are shown in stick with their molecular distance. (**B)**. 28 combinations of 8 different protein states of XylE WT and three mutants (D27N, E206Q and E206Q&D27N) in the presence or absence of the substrate (xylose) and 3 combinations of XylE WT and mutant D27N in the presence or absence of inhibitor (glucose) were studied in this work. The structures were generated by PyMol. The mutated residues are indicated by a star.

### Protonation of D27 controls the conformational transition

We first set out to understand the effect of protonation on the dynamics of XylE in the absence of substrate or inhibitor. To this end, we carried out ΔHDX-MS experiments comparing the WT protein with the mutants. We observe that protonation mimics D27N and E206Q cause an overall decrease in deuterium uptake on both the extracellular and intracellular sides compared to the WT protein (**Fig. 2A**). No significant exchange is observed in the transmembrane regions, which are mostly solvent inaccessible. Interestingly, the double mutant E206Q&D27N shows a decrease of deuterium uptake on the extracellular side coupled to an increase on the intracellular side - this corresponds to a ΔHDX pattern typical for the transition of transporter towards an IF state (**Fig. 2B**) [11]. To understand the sequence of events enabling protein transition to the IF state, we carried out ΔHDX-MS experiments comparing the single to double mutants. By comparing the double mutant E206Q&D27N to single mutant E206Q, we found ΔHDX pattern typical of a transition toward an IF state (**Fig. 2C**). By contrast the ΔHDX of E206Q&D27N versus D27N only showed minor differences in deuterium uptake (**Fig. 2D and Fig S3**). Taken together, these results suggest that D27 protonation is the main driver of the conformational transition to IF state. To confirm that the ΔHDX observed in our experiments was the result of conformational changes and not changes in global stability caused by the mutations we performed thermal unfolding experiments, monitored by circular dichroism (CD) measurements under temperature gradient [27, 28]. No significant change in stability was observed between the WT and the mutants below 50 °C (**Fig.S5**), which comforted us that the changes observed with HDX were mainly caused by conformational changes.

**Figure 2.**
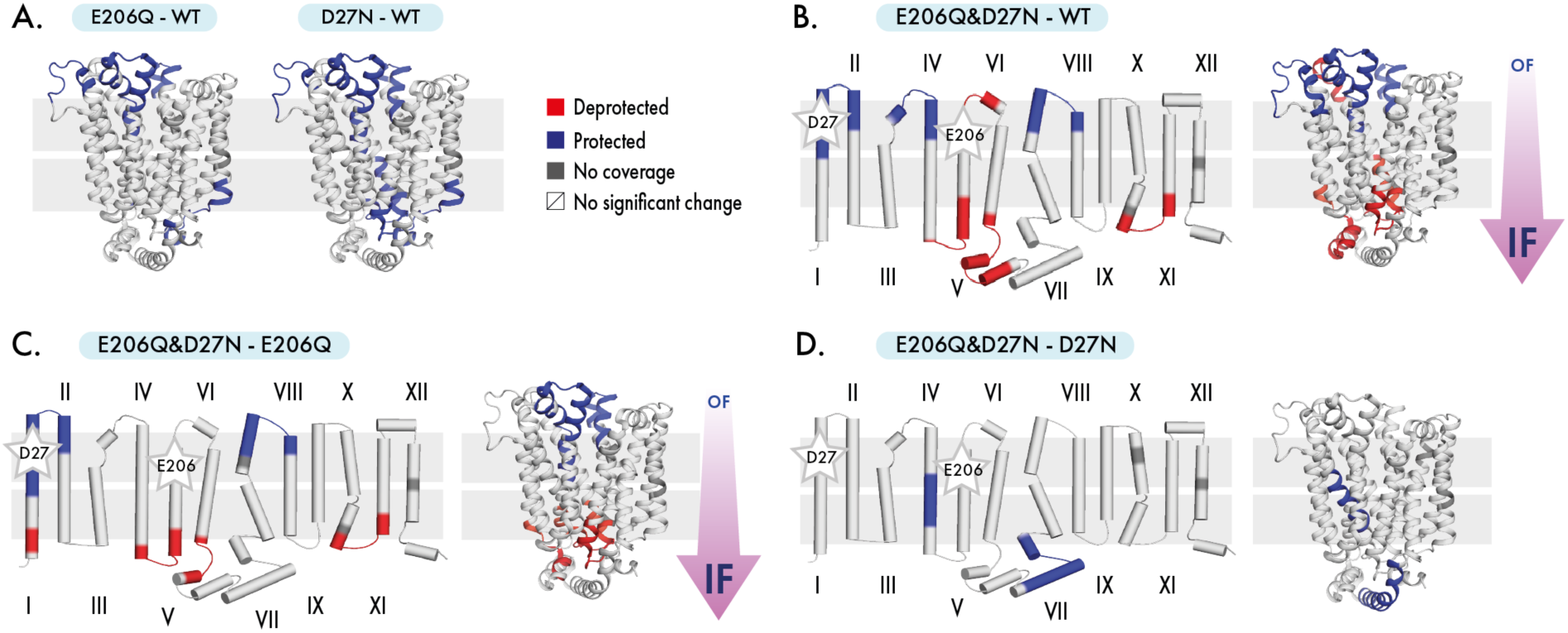
Conformational change through mutation (protonation mimic). **(A)**. Differential uptake pattern (ΔHDX) map between XylE E206Q or D27N and apo state. (**B)**. ΔHDX map between XylE E206Q&D27N and apo state. **C**. ΔHDX map between XylE E206Q&D27N and XylE D27N. Figures are plotted onto topological and 3D protein structure (PDB: 4GBY) by PyMol. Blue and red regions suggest a negative (protected) and a positive (deprotected) deuterium uptake pattern, respectively. The mutated residues are indicated by a star.

### Substrate and inhibitor binding favours the OF state

We then investigated the role of the substrate xylose and inhibitor glucose on the conformational equilibrium of XylE. The protein and mutants were incubated with 750 µM of xylose and the effect was followed by HDX-MS. The comparison between the protein in the presence and absence of xylose consistently shows that the presence of the substrate leads to a ΔHDX pattern typical of a transition towards an outward-open conformation; an increase in deuterium uptake on the extracellular side coupled to a decrease in deuterium uptake on the extracellular side (**Fig. 3A**). We performed similar experiments with the inhibitor glucose (750 µM). We observed that glucose also stabilizes the OF conformation regardless of the presence of mutations (**Fig. 3B**). Taken together, these results indicate that xylose or glucose binding induces a similar structural rearrangement towards the OF conformation regardless of the initial conformational state of the transporter, a finding in line with the OF states of the ligand-bound structures captured by X-ray crystallography [18]. However, this raises the question about how the transition of the loaded transporter toward the IF conformation occurs.

**Figure 3.**
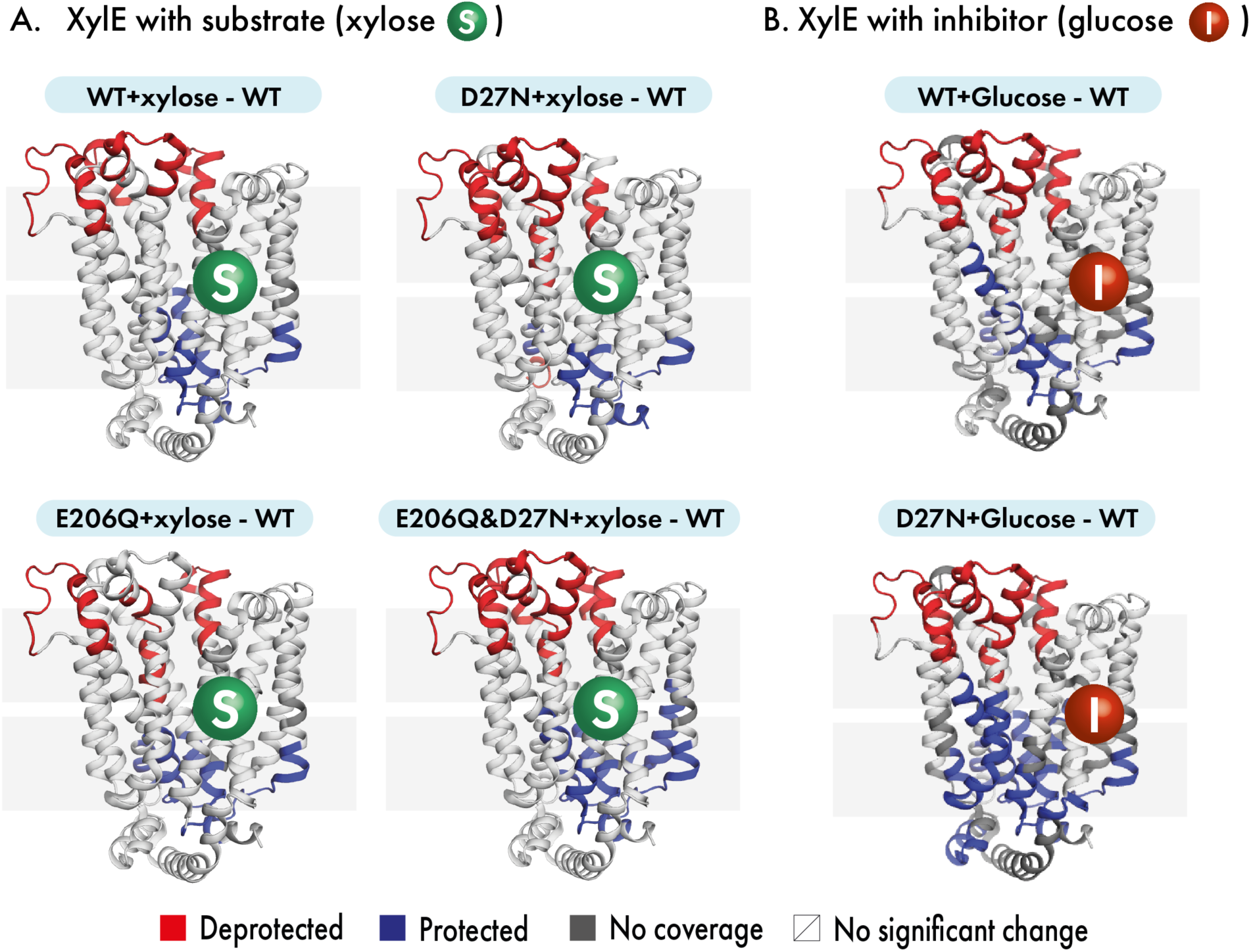
Conformational change through substrate/inhibitor binding. **(A)**. Differential deuterium uptake pattern between substrate (xylose) bound state with apo state. (**B)**. Differential deuterium uptake pattern between inhibitor (glucose) bound state with apo state. Figures are plotted onto 3D protein structure (PDB: 4GBY). Blue and red regions suggest a relatively negative (protected) and a positive (deprotected) deuterium uptake pattern respectively.

### Allosteric coupling between D27 protonation and substrate binding

Next, we went on to characterize how the combined effect of substrate binding and protonation mimics impacts on the conformational dynamics, to emulate a fully loaded transporter. We carried out ΔHDX-MS experiments of the mutant proteins versus the WT, in the presence of xylose. Strikingly, we observed that D27N versus WT in the presence of xylose (**Fig. 4A**) presented a different ΔHDX pattern compared to the apo experiment (**Fig. 2A**). The mutation leads to an increase in deuterium uptake on both sides of the protein, a pattern different from all the other ΔHDX patterns observed so far. The increased uptake on both sides of the transporter indicate that there is an increase in local structural dynamics on the entire protein. We hypothesize that the combined presence of the mutation and the substrate leads to increased conformational heterogeneity and label this new ΔHDX pattern as the “active state”. A similar ΔHDX pattern was observed when comparing D27N minus E206Q, the double mutant E206Q&D27N minus E206Q, but not E206Q minus WT, suggesting that the coupling is specific to D27 (**Fig. 4B, C, D**). We then performed the same experiment comparing D27N with the WT in the presence of the inhibitor glucose. To our surprise, this time we observed a pattern consistent with an OF conformation, suggesting that glucose binding tips the conformational equilibrium even more towards the OF state. This indicates that only a *bona fide* substrate can lead to the active state (**Fig. 4C**). Overall, our results suggest that D27 protonation is the trigger for activation of the protein while protonation of E206 has little effect. The shift toward this active state clearly demonstrates that an allosteric interplay exists between the mutation/protonation of D27 and binding of the substrate xylose.

**Figure 4.**
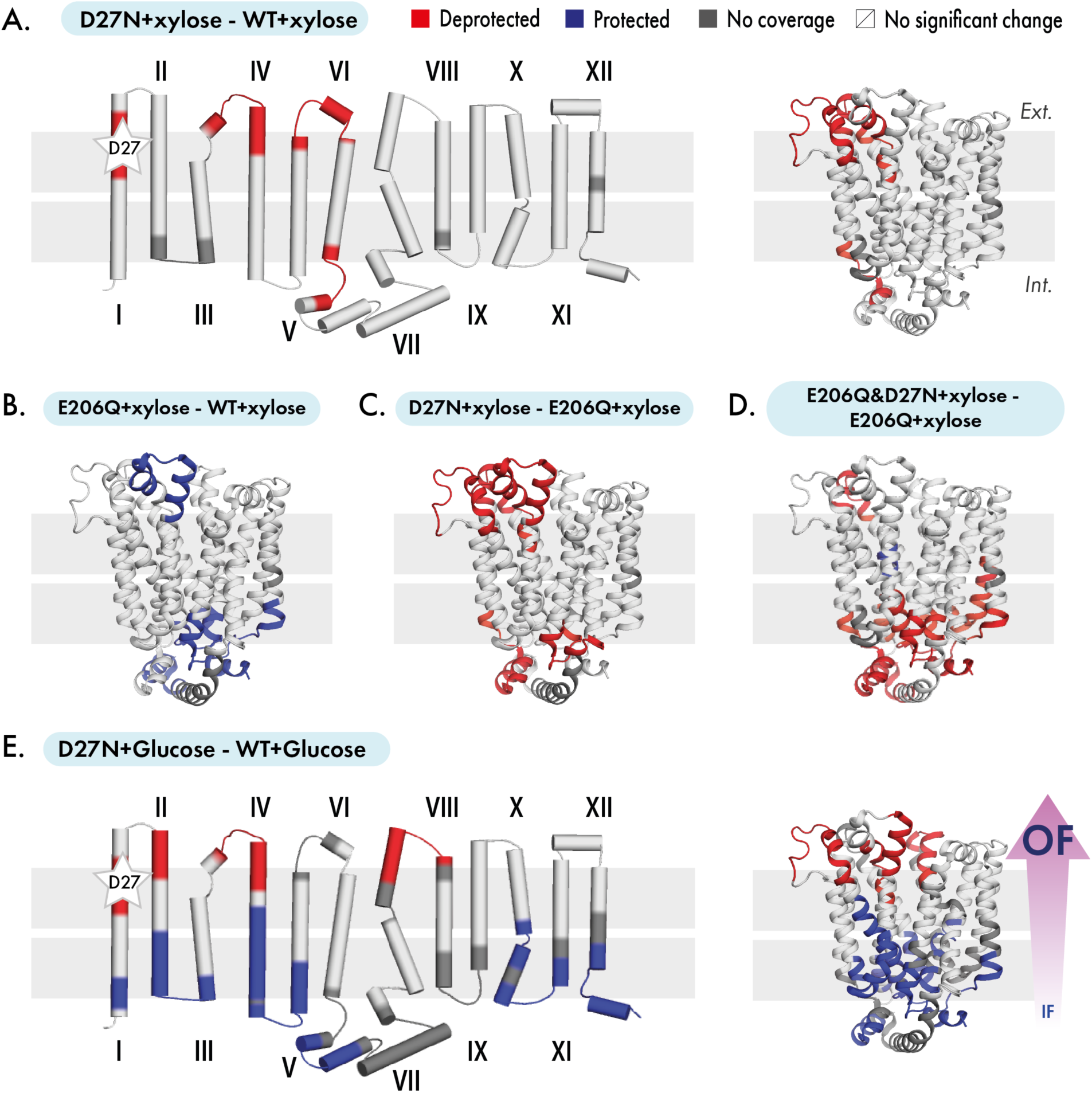
Conformational change through synergistic effect of substrate/inhibitor binding and mutation. **(A)**. ΔHDX map between XylE D27N with XylE wild type in the presence of substrate (xylose). (**B)**. ΔHDX map between XylE E206Q and XylE wild type in the presence of substrate (xylose). **(C)**. ΔHDX map between XylE D27N and XylE E206Q in the presence of substrate (xylose). (**D)**. ΔHDX map between XylE E206Q&D27N and XylE E206Q in the presence of substrate (xylose). (**E)**. ΔHDX map between XylE D27N with XylE wild type in the presence of inhibitor (glucose). Figures are plotted onto topological and 3D protein structure (PDB: 4GBY) by PyMol. Blue and red regions suggest a relatively negative (protected) and a positive (deprotected) deuterium uptake pattern respectively. The mutated residues are indicated by a star.

### MD simulations suggest double protonation leads to active state

In order to confirm the allosteric interplay between D27 and xylose binding, we ran all-atoms molecular dynamics (MD) simulations on the ligand-bound structures to visualize the effect of the protonation states of D27 on the protein structural dynamics. We calculated the intrinsic pK_a_ values of the residues D27 and E206 in the crystal structure using PROPKA [29]. The pK_a_ of D27 is 4.35 and that of E206 is 12.13. The intrinsic pK_a_ values of these residues suggests that the conformation captured in the crystal structure is protonated at E206 and deprotonated at D27.

We performed MD simulations of XylE embedded in a POPE lipid bilayer with the residue D27 either unprotonated or protonated, and E206 always protonated, using either the xylose-bound or glucose-bound structure. We clearly observe that in the case of unprotonated D27, xylose remains stably bound, essentially retaining the crystal structure pose (**Fig. 5a, d**). In contrast, xylose adopts multiple rotameric states in D27 protonated state (**Fig. 5b, e)**, suggesting that xylose binding stability is conditional on the presence of one proton on D27. Furthermore, instability of xylose is facilitated by the increased solvation of the substrate binding site (**Fig. 5g, h**). In contrast, the glucose bound simulation with D27 protonated retains the crystallographic pose with essentially a similar pattern of substrate stability and solvation as the xylose bound D27 unprotonated state (**Fig.5 c, f, i**).

**Figure 5.**
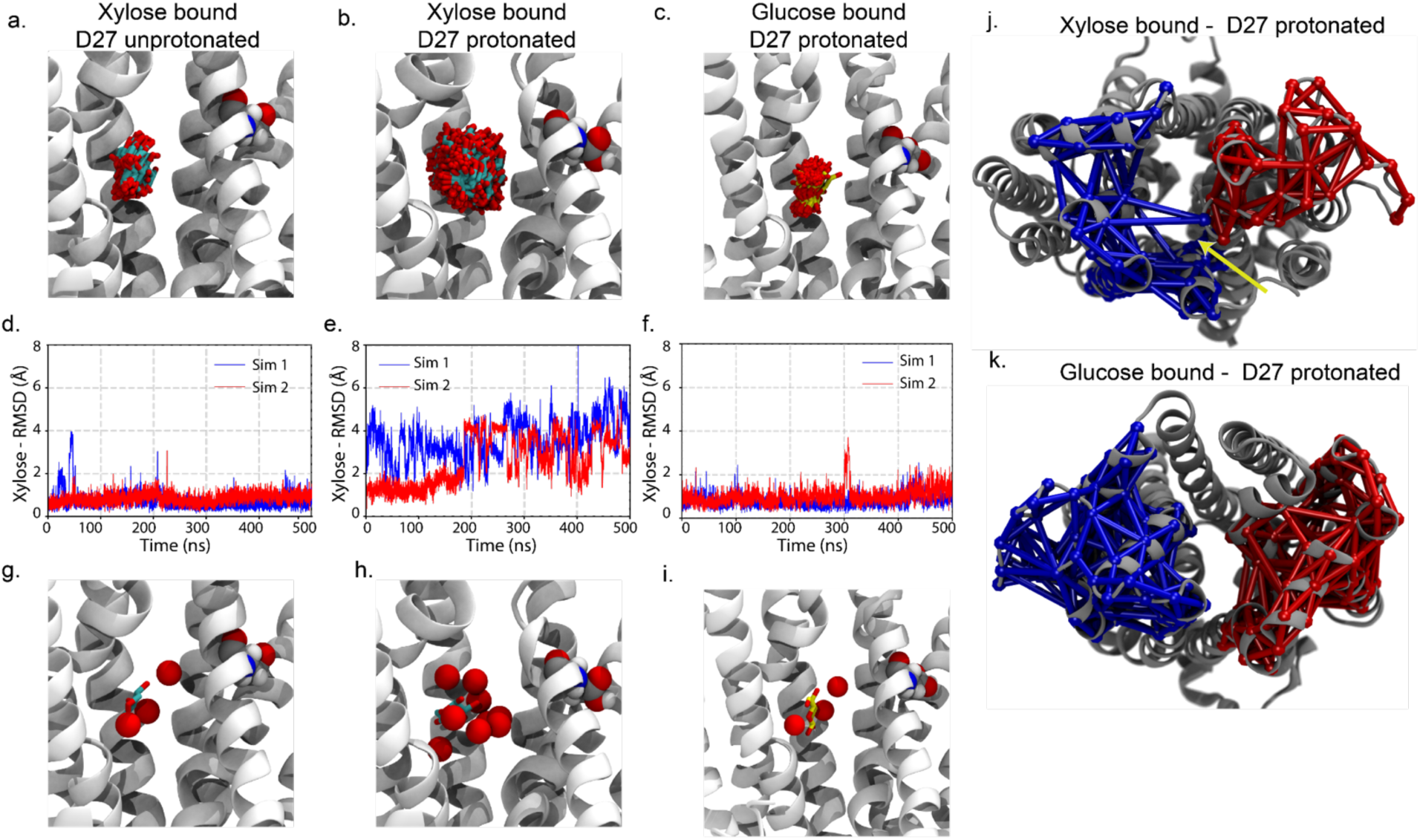
MD simulation reveal differential dynamics of XylE in substrate and inhibitor bound state. (**a**) MD simulations of xylose (shown in cyan and red sticks) bound XylE in the D27 unprotonated state, highlights that the bound substrate remains stable through the two independent 500 ns long MD simulations. (**b**) MD simulations of xylose and (**c**) glucose bound to protonated D27. The stability was characterized by monitoring the (**d-f**) RMSD of the xylose or glucose with respect to the crystal structure pose. (**g-i**) Solvation of the substrate binding site (defined by a sphere of radius 5 Å around xylose). Bound glucose molecule (shown in yellow and red sticks) in D27 protonated XylE retains the crystallographic pose with a similar pattern of substrate stability and solvation as the xylose bound unprotonated state. Allosteric coupling in extracellular gate of: (**j**) Xylose-bound and (**k**) Glucose-bound XylE in protonated state. Two distinct communities (shown in red and blue) captured from our 500 ns long MD simulations highlight correlated motions of the extracellular gate in the xylose-bound conformation. While, in glucose-bound conformation the extracellular gates remained de-coupled.

Dynamic network analysis was employed to better understand the differential conformational dynamics of XyIE in the presence of bound xylose and glucose. We captured correlated motions of the extracellular gate in the presence of bound xylose (**Fig. 5j**), while in the presence of bound glucose no such correlated motions of the extracellular gate were captured, and the transporter predominantly exists in the OF conformation (**Fig. 5k**).

The MD predictions corroborate the HDX-MS results at several levels. First, the high calculated pK_a_ value of E206 suggests that this residue is protonated most of the time during HDX-MS experiments carried out at pH7.4. This explains why E206Q mutation leads to minor or no changes in ΔHDX-MS experiments. Second, the instability of xylose binding and the increase of water molecules along the substrate pathway observed upon D27 protonation matches with the global increase in H/D exchange observed (**Fig.4 a, c, d**). Third, the correlated motions of the extracellular gates observed only in the presence of xylose confirms the existence of an active state with a greater conformational heterogeneity. The combination of MD predictions and HDX-MS results suggests that the combined presence of xylose and a proton on D27 leads to an unstable state that allows the conformational transition underlying transport.

### Discussion – revisiting the transport cycle

As a symporter, XylE binds and co-transport protons alongside its substrate xylose. Whether xylose or the proton needs to bind first is difficult to measure conclusively. Different studies have shown that XylE is protonated prior to substrate binding [19, 30] and hypothesized that D27 is the proton binding residue [20, 31]. However, our HDX experiments have singled out D27 as the driver of the conformational transition, but only if xylose is already present. We have also shown that substrate xylose and inhibitor glucose binding stabilize the outward-open conformation. These findings indicate that the initial protonation step is likely to happen on another residue than D27 and we propose that E206 plays this role. The most striking result of this work shows that D27N leads to an active state only if xylose is already bound, highlighting an allosteric coupling between the two residues. This effect is specific to xylose and shows that the protein can distinguish between substrate and inhibitor.

We propose the following transport cycle: in its resting state, the WT transporter is protonated at residue E206 most or all of the time. Binding of xylose to the protonated transporter stabilizes the OF conformation and facilitates solvent accessibility to residue D27. We speculate that the protonation of D27 occurs through water contact (**Fig. 6)**. The protonation of D27 when xylose is bound leads to a high energy activated state which initiates the conformational switch. This activated state is accessible only through allosteric coupling between D27 and the substrate binding site, and such coupling is exquisitely sensitive to xylose binding. Under transport conditions (e.g. in the presence of a proton gradient), XylE can then switch toward the inward facing conformation and release substrate and proton in the cytosol. According to the crystal structures, D27 is not exposed directly to the solvent in the inward open conformation and we speculate that a network of water molecules and acidic residues located on the cytoplasmic face – such as the highly conserved motif A – plays the role of proton transfer network [20]. Our study allows us to update the current knowledge of the transport mechanism of the prototypical secondary transporter XylE. The identification of D27 as the driver of the conformational transition correlates with the known role of equivalent residues for others proton coupled MFS transporters such as LacY (E325), LmrP (E327), MdfA (D34) and YajR (E320) [32]. This suggests a conserved mechanism of action among proton coupled symporters of the same structural family. We surmise that, along the resolution revolution, the development of tools and workflows capable of answering mechanistic questions at a molecular level is much-needed, and we demonstrate that HDX-MS coupled to MD simulations have a key role to play.

**Figure 6.**
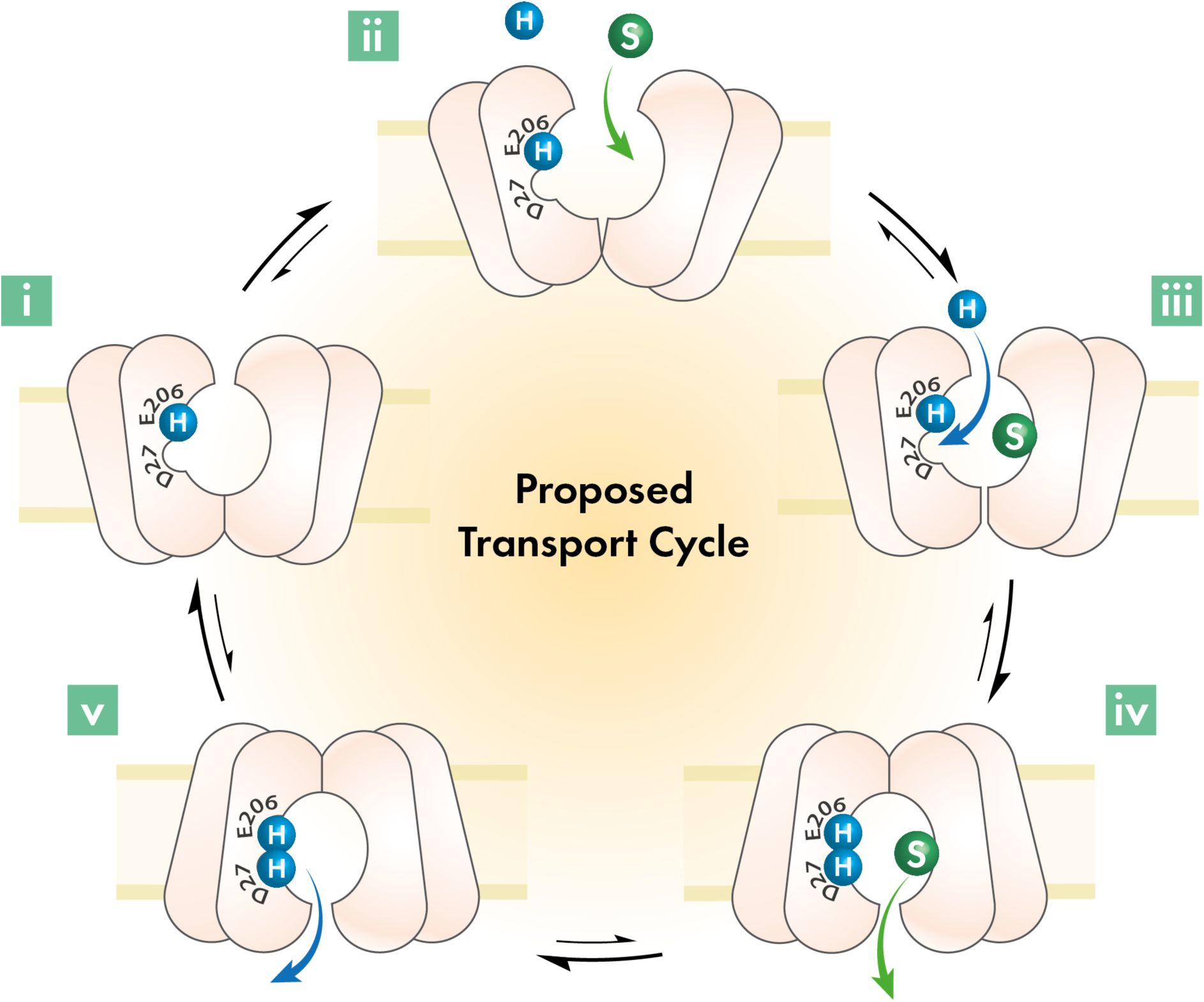
Proposed model of XylE active transport. Model was deduced through combinations of differential HDX-MS experiments designed to mimic the transport cycle of XylE. **(i)** E206 is always protonated. **(ii)** Substrate binding. **(iii)** Second proton binds to D27, destabilizing the protein. **(iv)** Substrate release. **(v)** Proton release and it goes to next round of switch. Proton are indicated as blue sphere with an “H”, substrate is labeled as an “S”.

## Methods

### XylE expression and purification

XylE was overexpressed in *E. coli* BL21-AI (DE3) (Invitrogen), which was transformed with the XylE gene in the presence or absence of the chosen mutations and cloned in the (30 µg/ml) kanamycin-resistant pET28-a plasmid (Novagen) modified with a C-terminal 10-histidine tag, grown in 6 baffled flasks each containing 1 L of LB media at 37 °C 220 r.p.m to an OD_600_ of 0.8. Expression was induced with 1 mM IPTG (Isopropy-b-D-1-thiogalactopyranoside) and 0.1% (w/v) L-arabinose and growth continued until the value of OD_600_ is flat. The cells were harvested by centrifugation, washed in 200 mL PBS buffer and centrifuged again for 20 min at 4,200 r.p.m in a Beckman JLA-16.250 rotor. The pellet was then resuspended in 50 mL PBS (phosphate-buffered saline) with 10 mM β-mercaptoethanol and 1 cOmplete protease inhibitor tablet and was frozen at −70 °C before purification. Cells were defrosted and incubated with 1.5 µL benzonase nuclease (ThermoFisher) for 10 min at room temperature before passed through constant cell disrupter at 25 kPsi, 4 °C. Then the ice-chilled membranes were isolated by ultracentrifugation for 30 min at 38,000 r.p.m in a Beckman Ti45 rotor, 4 °C. Membrane pellets were solubilised for 2 hours with mixing in solubilisation buffer [50 mM sodium phosphate pH 7.4, 200 mM NaCl, 10% (v/v) glycerol, 20 mM imidazole, 10 mM β-mercaptoethanol, and 2% n-Dodecyl β-D-maltoside (β-DDM, Anatrace), 0.1 mM phenylmethylsulfonyl fluoride (PMSF) and EDTA free protease inhibitor tablet (Roche)] at 4 °C. Then the protein solution was isolated by centrifugation for another 30 min at 38,000 r.p.m in a Beckman Ti70 rotor to remove DDM insoluble material. The supernatant was filtered using 0.45 µm filter and applied to a Ni-NTA column equilibrated in 96% SEC purification buffer [50 mM sodium phosphate pH 7.4, 10% (v/v) glycerol, 2 mM β-mercaptoethanol, and 0.05% β-DDM (Anatrace), 0.1 mM phenylmethylsulfonyl fluoride] and 4% elution buffer [50 mM sodium phosphate pH 7.4, 500 mM imidazole, 10% (v/v) glycerol, 10 mM β-mercaptoethanol, 0.1 mM phenylmethylsulfonyl fluoride (PMSF) and 0.05% β-DDM (Anatrace)]. The bound protein was washed with 50 mL 85% SEC purification buffer, 15% elution buffer and eluted with 2 mL of 100% elution buffer, which was collected for further size exclusion chromatography (SEC). The SEC purification was conducted with a Superdex 16/600 GL SEC column, which was equilibrated with SEC purification buffer. The elution fraction was collected and concentrated with a Vivaspin concentrator (100 kDa cutoff). The samples were either flash frozen and kept at −70 °C until use or used directly for HDX-MS experiments.

### Hydrogen-deuterium exchange mass spectrometry

Hydrogen-deuterium exchange mass spectrometry (HDX-MS) experiments were done as previously described [11] using a Synapt G2-Si HDMS coupled to nanoACQUITY UPLC with HDX Automation technology (Waters Corporation, Manchester, UK). Membrane proteins in detergent micelles were prepared at a concentration around 30 µM using a 100 kDa cutoff Vivaspin concentrators. Before quenching in 100 μL ice cold buffer Q (100mM potassium phosphate in formic acid pH 2.4), each 5 μL protein sample was incubated for 30 s, 5 min and 30 min in 95 μL deuterium labelling buffer L (10 mM potassium phosphate in D_2_O pH 6.6). Then, the protein was digested with self-packed pepsin column at 20 °C. The reference controls were performed using the same protocol, while incubated with 95 μL equilibration buffer E (10 mM potassium phosphate in H_2_O pH 7.0) instead. The pepsin column was washed between injections using pepsin wash buffer (1.5 M Gu-HCl, 4% (v/v) MeOH, 0.8% (v/v) formic acid). A clean blank run was done between each sample run to reduce peptide carry-over. Peptides were trapped for 3 minusing an Acquity BEH C18 1.7 μM VABGUARD pre-column at a 200 μL/min flow rate in buffer A (0.1% formic acid in HPLC water pH 2.5) before eluted to an Acquity UPLC BEH C18 1.7 μM analytical column with a linear gradient buffer B (8-40% gradient of 0.1% formic acid in acetonitrile) at a flow rate of 40 μL/min. Then peptides went through electrospray ionization progress in a positive ion mode using Synapt G2-Si mass spectrometer (Waters). Leucine Enkephalin was applied for mass accuracy correction and sodium iodide was used as calibration for the mass spectrometer. MS^E^ data was collected by a 20-30 V trap collision energy ramp. All the isotope labelling time points were performed in triplicates.

### HDX data evaluation and statistical analysis

Acquired reference MS^E^ data was put on PLGS (ProteinLynx Global Server 2.5.1, Waters) to achieve the peptide map, then all the MS^E^ data including reference and deuterated samples were processed by DynamX v.3.0 (Waters) for deuterium uptake determination. Peptide filtration and analysis were performed as described before [11]. Woods plots were generated by the in-house Deuteros software [26].

### Circular dichroism measurements

CD thermal denaturation was performed in an Aviv Circular Dichroism Spectrophotometer, Model 410 (Biomedical Inc., Lakewood, NJ, USA). All the samples of XylE were measured at a protien concentration at 0.14 – 0.17 mg.ml^-1^ scale range and a cell pathlength of 1mm. The sample was heated at 5 °C intervals in SEC (size exclusion Chromatography) purification buffer (50 mM sodium phosphate, 10% (v/v) glycerol, 2 mM β-mercaptoethanol, and 0.05% β-DDM (Anatrace), 0.1 mM phenylmethylsulfonyl fluoride, pH 7.4) from 25 °C to 95 °C. Each sample was scanned two times at fixed wavelength 222 nm in 1 nm wavelength step with averaging time of 1 s. The mean residue ellipticity at 222 nm was used for further analysis. The calculation of mean residue ellipticity ([θ]_mrw_) is following the equation:

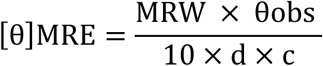

Where, θ_obs_=observed ellipticity in degrees, d = pathlength in cm, c = concentration in mg/ml. Mean residue weight (MRW) is calculate from the molecular mass divided by the number of amino acids - 1 and the value of MRW is around 110 for most of the proteins.

### Molecular dynamics: Simulation Setup

Molecular dynamics simulations were initiated from either xylose (PDB ID: 4GBY) or glucose bound (PDB ID: 4GBZ) state of XylE [18]. Protonation states of the titratable residues were assigned based on pKa calculations performed using PROPKA3.1 at pH 7 [29]. Thereafter, XylE was embedded in a POPE lipid bilayer using the membrane replacement method in CHARMM-GUI [33]. System was solvated with TIP3P water molecules [34]. Thereafter, Na^+^ and Cl^−^ ions were added, and the system was neutralized with the ionic concentration set to 100 mM. The final system inclusive of the protein, lipids, water molecules, and ions comprised ∼100 K atoms.

Subsequently, the system was relaxed by minimizing it to a minimum for 5000 steps using conjugate-gradient algorithm and simulated for 5 ns at 310 K, with all the heavy atoms of the protein and the substrate restrained to their crystallographic positions with a force constant of k = 5 kcal/mol/Å^2^. Finally, all the restrains were removed and the systems were simulated for 500 ns.

### MD simulation protocol

The simulations were performed on with NAMD 2.13 [35] employing CHARMM36 protein and lipid forcefields [36]. Simulations were performed in an NPT ensemble with periodic boundary conditions. Temperature was maintained at 310 K using Langevin dynamics with a damping constant of 0.5 ps^-1^. Pressure was maintained at 1 atm using the Nosé–Hoover Langevin piston method [37]. The cutoff used for the short range interactions were 12 Å with the switching applied at 10 Å. Long-range electrostatics was treated by the employing particle mesh Ewald (PME) algorithm [38]. Bonded, non-bonded, and PME calculations were performed at 2-, 2-, and 4-fs intervals, respectively.

### Analysis: Dynamical network analysis

In XylE, coupling in the extracellular and intracellular gates can be understood in terms of the allosteric interactions of residues that efficiently move in a correlated manner. For this, dynamic network analysis was performed using the Network-View plugin [39] in VMD. In a network, all Cα carbons are defined as nodes connected by edges if they are within 4.5 Å of each other for at least 75% of the MD trajectory. Pearson correlation was used to define the communities in the entire network corresponding to the set of residues that move in concert with each other.

## References

1. Cheng, Y., Membrane protein structural biology in the era of single particle cryo-EM. Curr Opin Struct Biol, 2018. 52: p. 58–63.

2. Moraes, I., et al., Membrane protein structure determination - the next generation. Biochim Biophys Acta, 2014. 1838 (1 Pt A): p. 78–87.

3. Forrest, L.R., R. Kramer, and C. Ziegler, The structural basis of secondary active transport mechanisms. Biochim Biophys Acta, 2011. 1807(2): p. 167–88.

4. Drew, D. and O. Boudker, Shared Molecular Mechanisms of Membrane Transporters. Annu Rev Biochem, 2016. 85: p. 543–72.

5. LeVine, M.V., et al., Allosteric Mechanisms of Molecular Machines at the Membrane: Transport by Sodium-Coupled Symporters. Chem Rev, 2016. 116(11): p. 6552–87.

6. Moradi, M. and E. Tajkhorshid, Mechanistic picture for conformational transition of a membrane transporter at atomic resolution. Proc Natl Acad Sci U S A, 2013. 110(47): p. 18916–21.

7. Konermann, L., J. Pan, and Y.H. Liu, Hydrogen exchange mass spectrometry for studying protein structure and dynamics. Chem Soc Rev, 2011. 40(3): p. 1224–34.

8. Engen, J.R., Analysis of protein conformation and dynamics by hydrogen/deuterium exchange MS. Anal Chem, 2009. 81(19): p. 7870–5.

9. McHaourab, H.S., P.R. Steed, and K. Kazmier, Toward the fourth dimension of membrane protein structure: insight into dynamics from spin-labeling EPR spectroscopy. Structure, 2011. 19(11): p. 1549–61.

10. Liang, B. and L.K. Tamm, NMR as a tool to investigate the structure, dynamics and function of membrane proteins. Nat Struct Mol Biol, 2016. 23(6): p. 468–74.

11. Martens, C., et al., Direct protein-lipid interactions shape the conformational landscape of secondary transporters. Nat Commun, 2018. 9(1): p. 4151.

12. Reading, E., et al., Interrogating Membrane Protein Conformational Dynamics within Native Lipid Compositions. Angew Chem Int Ed Engl, 2017. 56(49): p. 15654–15657.

13. Skinner, J.J., et al., Protein dynamics viewed by hydrogen exchange. Protein Sci, 2012. 21(7): p. 996–1005.

14. Vadas, O. and J.E. Burke, Probing the dynamic regulation of peripheral membrane proteins using hydrogen deuterium exchange-MS (HDX-MS). Biochem Soc Trans, 2015. 43(5): p. 773–86.

15. Martens, C., et al., Integrating hydrogen-deuterium exchange mass spectrometry with molecular dynamics simulations to probe lipid-modulated conformational changes in membrane proteins. Nat Protoc, 2019. 14(11): p. 3183–3204.

16. Shi, Y., Common folds and transport mechanisms of secondary active transporters. Annu Rev Biophys, 2013. 42: p. 51–72.

17. Quistgaard, E.M., et al., Structural basis for substrate transport in the GLUT-homology family of monosaccharide transporters. Nat Struct Mol Biol, 2013. 20(6): p. 766–8.

18. Sun, L., et al., Crystal structure of a bacterial homologue of glucose transporters GLUT1-4. Nature, 2012. 490(7420): p. 361–6.

19. Madej, M.G., et al., Functional architecture of MFS D-glucose transporters. Proc Natl Acad Sci U S A, 2014. 111(7): p. E719–27.

20. Wisedchaisri, G., et al., Proton-coupled sugar transport in the prototypical major facilitator superfamily protein XylE. Nat Commun, 2014. 5: p. 4521.

21. Bazzone, A., et al., pH Regulation of Electrogenic Sugar/H+ Symport in MFS Sugar Permeases. PLoS One, 2016. 11(5): p. e0156392.

22. Lento, C., et al., Time-resolved ElectroSpray Ionization Hydrogen-deuterium Exchange Mass Spectrometry for Studying Protein Structure and Dynamics. J Vis Exp, 2017(122).

23. Bai, X., T.F. Moraes, and R.A.F. Reithmeier, Structural biology of solute carrier (SLC) membrane transport proteins. Mol Membr Biol, 2017. 34(1-2): p. 1–32.

24. Bazzone, A., et al., A Loose Relationship: Incomplete H(+)/Sugar Coupling in the MFS Sugar Transporter GlcP. Biophys J, 2017. 113(12): p. 2736–2749.

25. Masson, G.R., et al., Recommendations for performing, interpreting and reporting hydrogen deuterium exchange mass spectrometry (HDX-MS) experiments. Nat Methods, 2019. 16(7): p. 595–602.

26. Lau, A.M.C., et al., Deuteros: software for rapid analysis and visualization of data from differential hydrogen deuterium exchange-mass spectrometry. Bioinformatics, 2019. 35(17): p. 3171–3173.

27. Greenfield, N.J., Using circular dichroism collected as a function of temperature to determine the thermodynamics of protein unfolding and binding interactions. Nat Protoc, 2006. 1(6): p. 2527–35.

28. Harris, N.J., et al., Comparative stability of Major Facilitator Superfamily transport proteins. Eur Biophys J, 2017. 46(7): p. 655–663.

29. Olsson, M.H., et al., PROPKA3: Consistent Treatment of Internal and Surface Residues in Empirical pKa Predictions. J Chem Theory Comput, 2011. 7(2): p. 525–37.

30. Smirnova, I., et al., Role of protons in sugar binding to LacY. Proc Natl Acad Sci U S A, 2012. 109(42): p. 16835–40.

31. Ke, M., et al., Molecular determinants for the thermodynamic and functional divergence of uniporter GLUT1 and proton symporter XylE. PLoS Comput Biol, 2017. 13(6): p. e1005603.

32. Zhang, X.C., et al., Energy coupling mechanisms of MFS transporters. Protein Sci, 2015. 24(10): p. 1560–79.

33. Jo, S., et al., CHARMM-GUI Membrane Builder for mixed bilayers and its application to yeast membranes. Biophys J, 2009. 97(1): p. 50–8.

34. Harrach, M.F. and B. Drossel, Structure and dynamics of TIP3P, TIP4P, and TIP5P water near smooth and atomistic walls of different hydroaffinity. J Chem Phys, 2014. 140(17): p. 174501.

35. Phillips, J.C., et al., Scalable molecular dynamics with NAMD. J Comput Chem, 2005. 26(16): p. 1781–802.

36. Best, R.B., et al., Optimization of the additive CHARMM all-atom protein force field targeting improved sampling of the backbone phi, psi and side-chain chi(1) and chi(2) dihedral angles. J Chem Theory Comput, 2012. 8(9): p. 3257–3273.

37. Glenn J. Martyna, G.J., Tobias, D. J., & Klein, M. L, Constant pressure molecular dynamics algorithms. The Journal of Chemical Physics, 1994. 101(5).

38. Ulrich Essmann, L.P., Max L. Berkowitz, A smooth particle mesh Ewald method. The Journal of Chemical Physics 1995. 103 (19).

39. Sethi, A., et al., Dynamical networks in tRNA:protein complexes. Proc Natl Acad Sci U S A, 2009. 106(16): p. 6620–5.

