## Supplemental Figures for "Hydrogen-deuterium exchange mass spectrometry captures distinct dynamics upon substrate and inhibitor binding to a transporter"

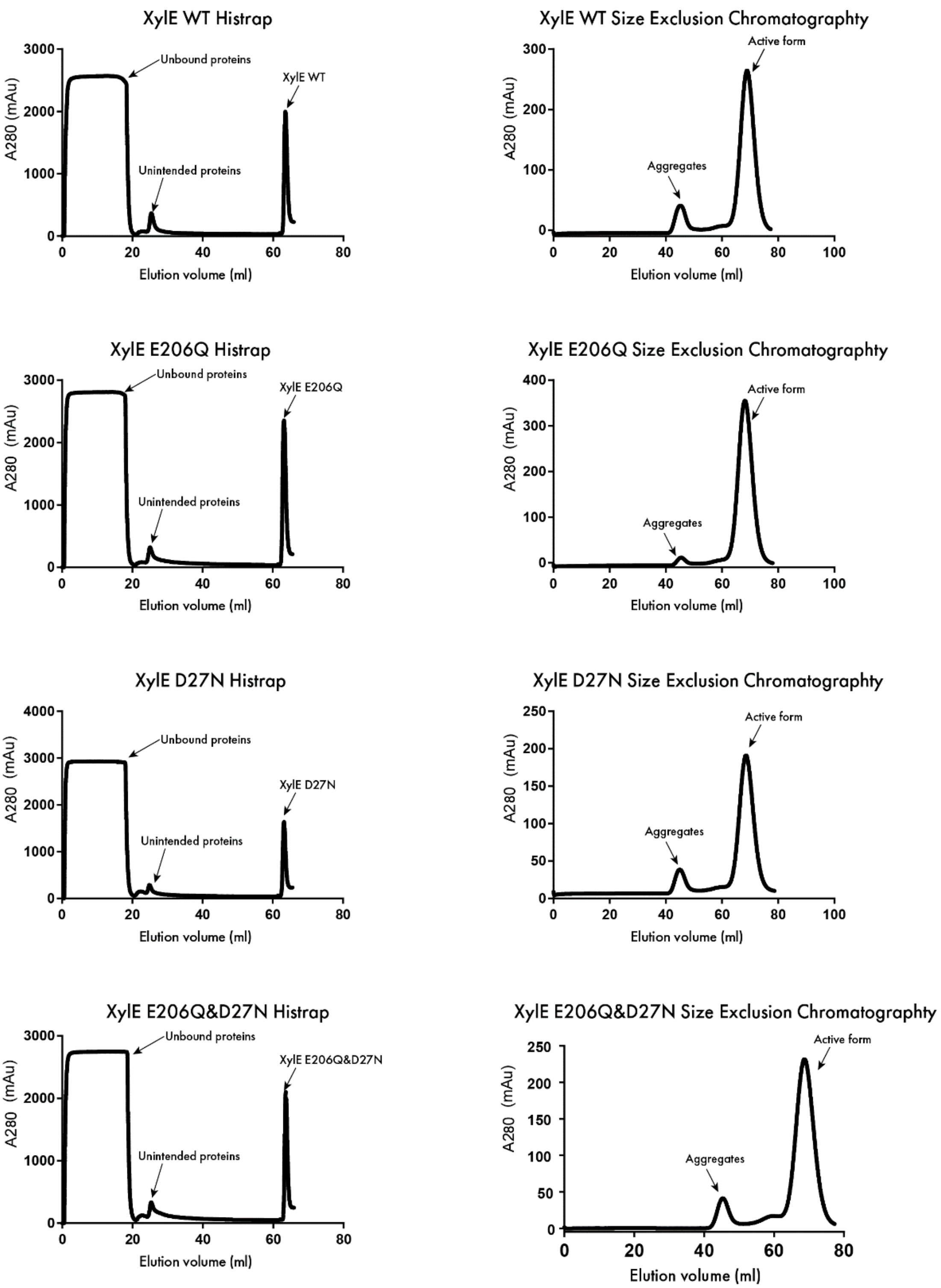


**Figure S1.** Histrap and Size Exclusion Chromatography (SEC) of XylE and associated mutants. The chromatography of Histrap and SEC were obtained by AKTA$^{TM}$ Pure

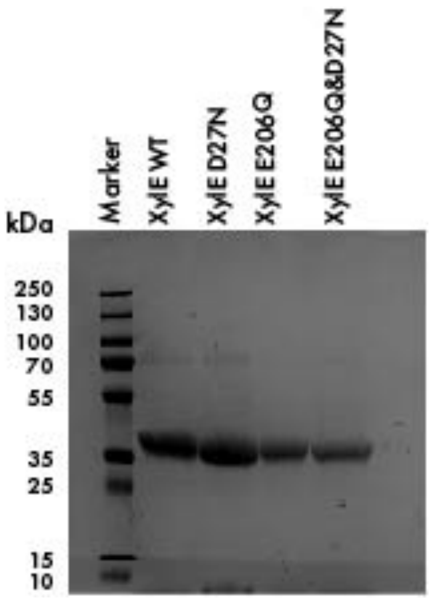


**Figure S2.** SDS-PAGE analysis of XylE wild type, D27N, E206Q, and E206Q&D27N.

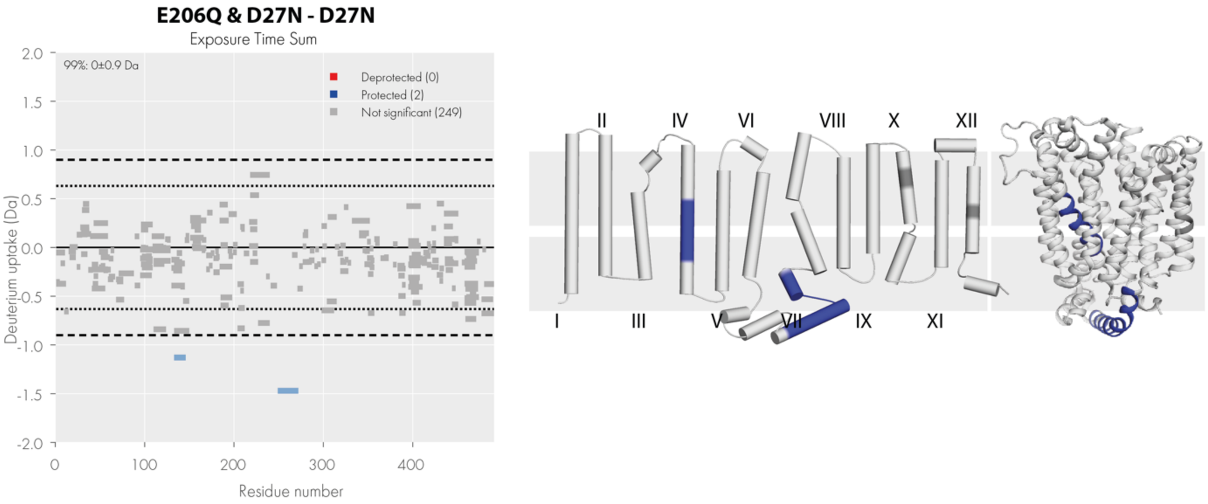


**Figure S3.** Woods plot of the combination between E206Q&D27N and D27N. Each bar represents a single peptide with peptide length indicated by the bar length. No major deuterium uptake difference can be seen from comparison. Figures are plotted onto topological and 3D protein structure (PDB: 4GBY) by PyMol. Blue and red regions suggest a relatively negative (protected) and a positive (deprotected) deuterium uptake pattern respectively.

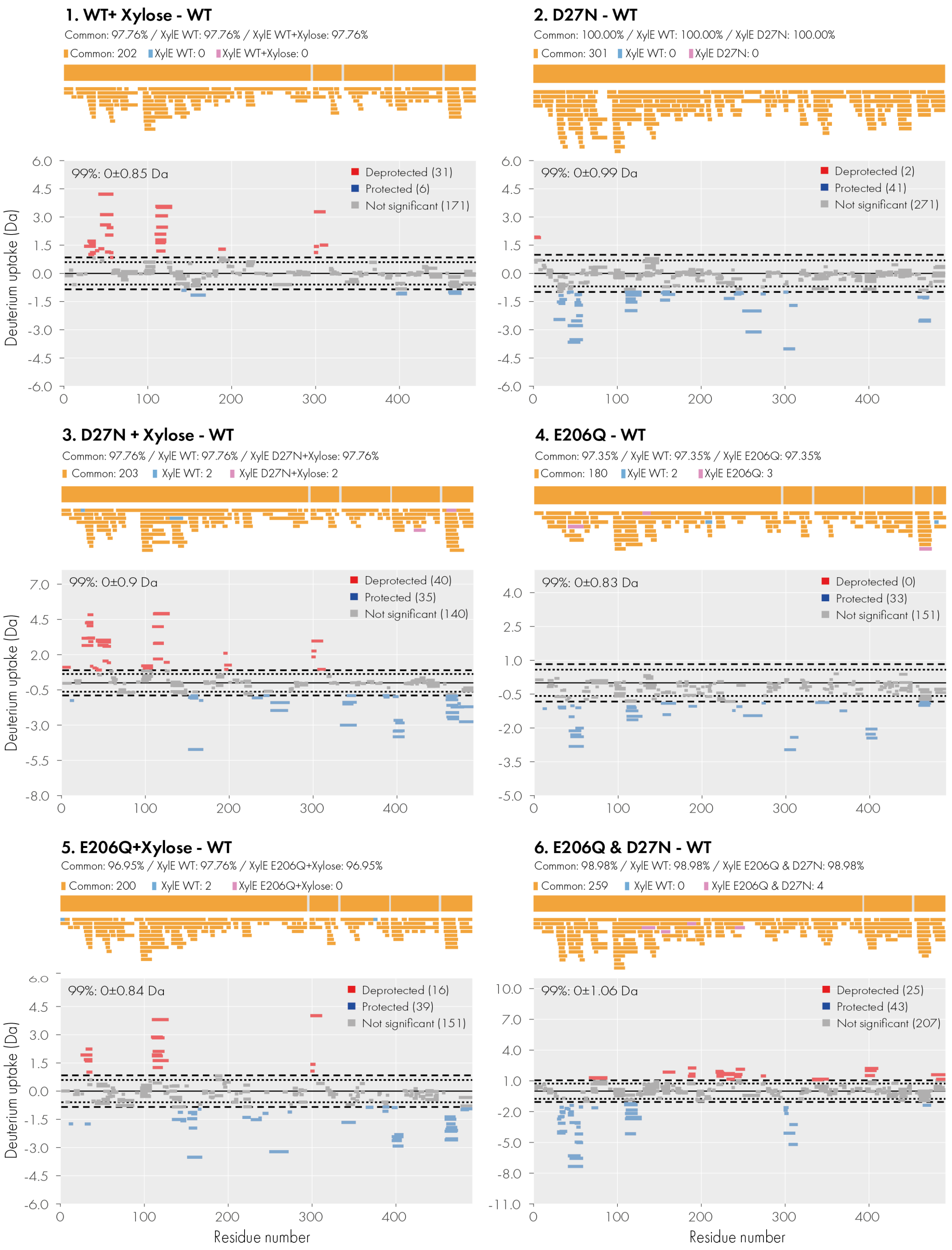
1. WT+ Xylose - WT
Common: 97.76% / XylE WT: 97.76% / XylE WT+Xylose: 97.76%
Common: 202
XylE WT: 0
XylE WT+Xylose: 0
99%: 0±0.85 Da
Deprotected (31)
Protected (6)
Not significant (171)
2. D27N - WT
Common: 100.00% / XylE WT: 100.00% / XylE D27N: 100.00%
Common: 301
XylE WT: 0
XylE D27N: 0
99%: 0±0.99 Da
Deprotected (2)
Protected (41)
Not significant (271)
3. D27N + Xylose - WT
Common: 97.76% / XylE WT: 97.76% / XylE D27N+Xylose: 97.76%
Common: 203
XylE WT: 2
XylE D27N+Xylose: 2
99%: 0±0.9 Da
Deprotected (40)
Protected (35)
Not significant (140)
4. E206Q - WT
Common: 97.35% / XylE WT: 97.35% / XylE E206Q: 97.35%
Common: 180
XylE WT: 2
XylE E206Q: 3
99%: 0±0.83 Da
Deprotected (0)
Protected (33)
Not significant (151)
5. E206Q+Xylose - WT
Common: 96.95% / XylE WT: 97.76% / XylE E206Q+Xylose: 96.95%
Common: 200
XylE WT: 2
XylE E206Q+Xylose: 0
99%: 0±0.84 Da
Deprotected (16)
Protected (39)
Not significant (151)
6. E206Q & D27N - WT
Common: 98.98% / XylE WT: 98.98% / XylE E206Q & D27N: 98.98%
Common: 259
XylE WT: 0
XylE E206Q & D27N: 4
99%: 0±1.06 Da
Deprotected (25)
Protected (43)
Not significant (207)
Deuterium uptake (Da)
Residue number

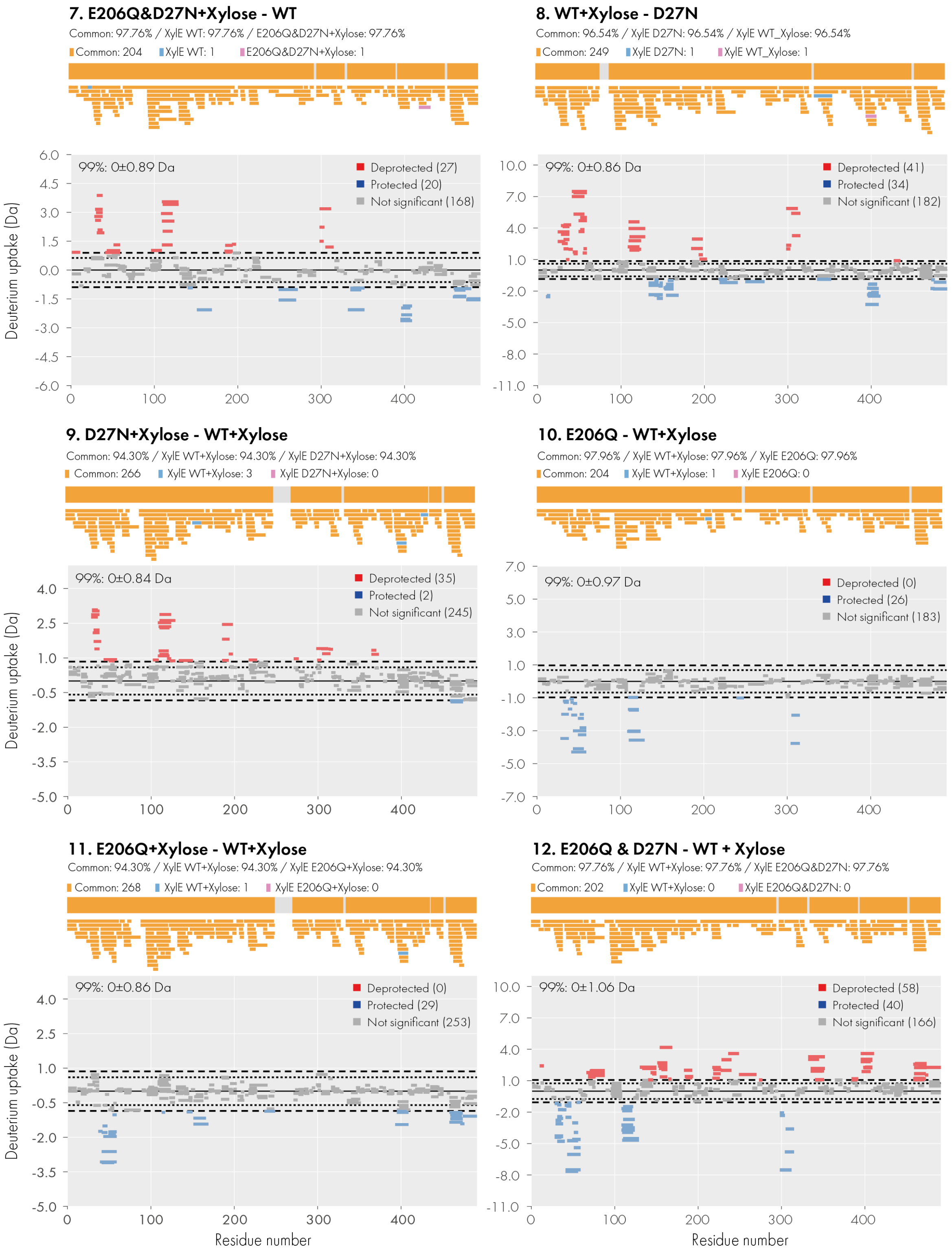
7. E206Q&D27N+Xylose - WT
Common: 97.76% / XylE WT: 97.76% / E206Q&D27N+Xylose: 97.76%
Common: 204
XylE WT: 1
E206Q&D27N+Xylose: 1
99%: 0±0.89 Da
Deprotected (27)
Protected (20)
Not significant (168)
8. WT+Xylose - D27N
Common: 96.54% / XylE D27N: 96.54% / XylE WT_Xylose: 96.54%
Common: 249
XylE D27N: 1
XylE WT_Xylose: 1
99%: 0±0.86 Da
Deprotected (41)
Protected (34)
Not significant (182)
9. D27N+Xylose - WT+Xylose
Common: 94.30% / XylE WT+Xylose: 94.30% / XylE D27N+Xylose: 94.30%
Common: 266
XylE WT+Xylose: 3
XylE D27N+Xylose: 0
99%: 0±0.84 Da
Deprotected (35)
Protected (2)
Not significant (245)
10. E206Q - WT+Xylose
Common: 97.96% / XylE WT+Xylose: 97.96% / XylE E206Q: 97.96%
Common: 204
XylE WT+Xylose: 1
XylE E206Q: 0
99%: 0±0.97 Da
Deprotected (0)
Protected (26)
Not significant (183)
11. E206Q+Xylose - WT+Xylose
Common: 94.30% / XylE WT+Xylose: 94.30% / XylE E206Q+Xylose: 94.30%
Common: 268
XylE WT+Xylose: 1
XylE E206Q+Xylose: 0
99%: 0±0.86 Da
Deprotected (0)
Protected (29)
Not significant (253)
12. E206Q & D27N - WT + Xylose
Common: 97.76% / XylE WT+Xylose: 97.76% / XylE E206Q&D27N: 97.76%
Common: 202
XylE WT+Xylose: 0
XylE E206Q&D27N: 0
99%: 0±1.06 Da
Deprotected (58)
Protected (40)
Not significant (166)
Deuterium uptake (Da)
Residue number

### 13. E206Q & D27N+Xylose - WT + Xylose

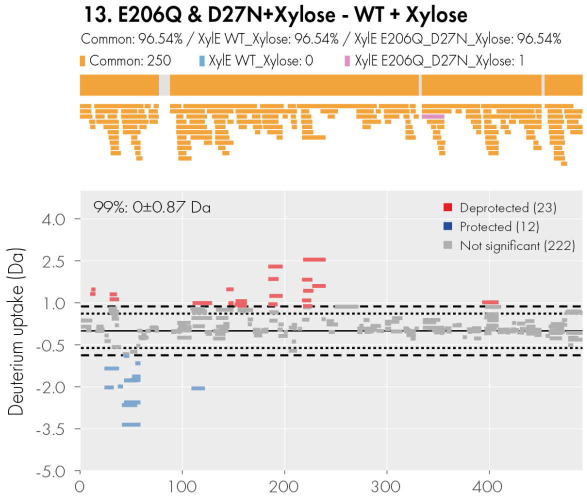


### 14. D27N + Xylose - D27N

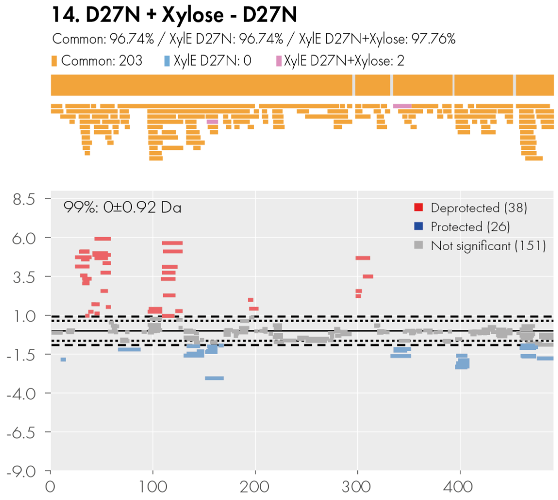


## 15. E206Q - D27N

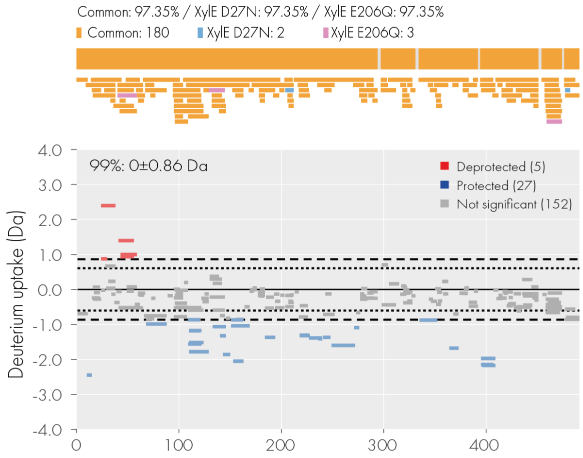


### 16. E206Q+Xylose - D27N

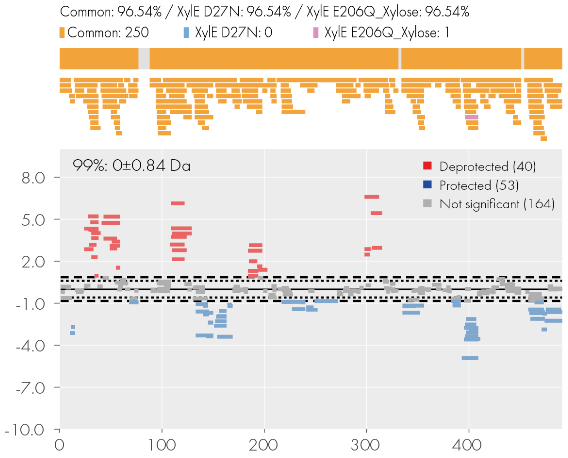


## 17. E206Q & D27N - D27N

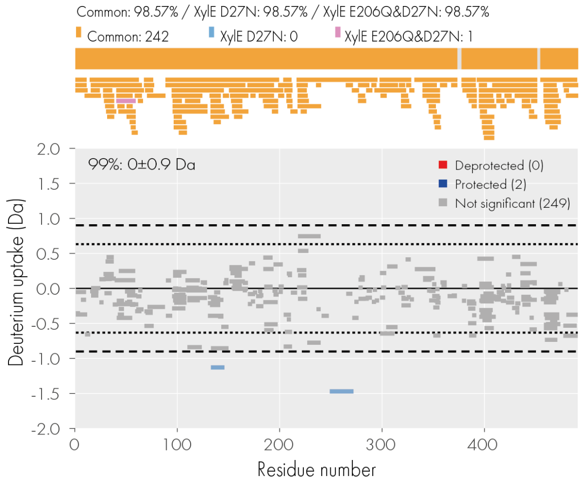


### 18. E206Q&D27N+Xylose - D27N

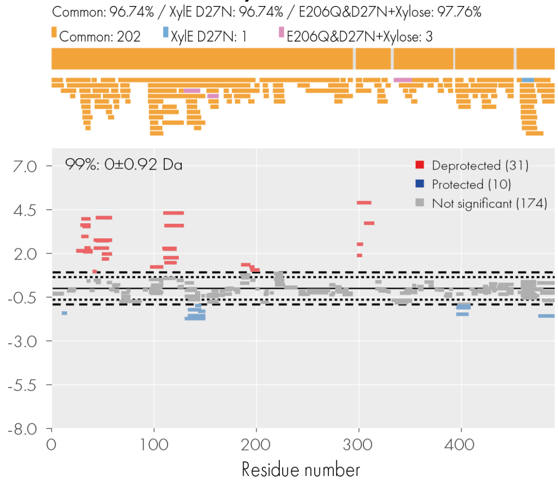

### 19. D27N + Xylose - E206Q

Common: 97.76% / XylE E206Q: 97.76% / XylE D27N+Xylose: 97.76%

Common: 202 XylE E206Q: 4 XylE D27N+Xylose: 3

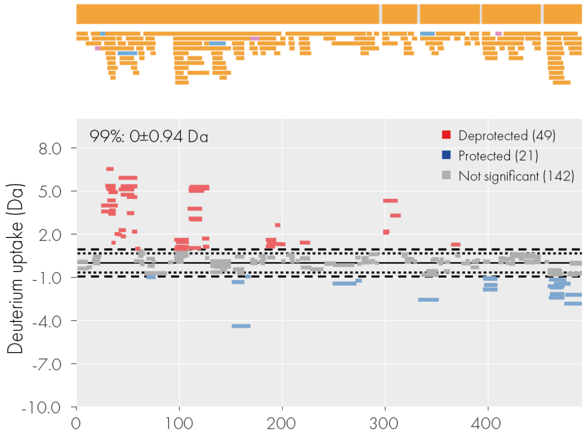


### 20. E206Q+Xylose - D27N+Xylose

Common: 93.89% / XylE D27N+Xylose: 93.89% / XylE E206Q+Xylose: 93.89%

Common: 247 XylE D27N+Xylose: 9 XylE E206Q+Xylose: 5

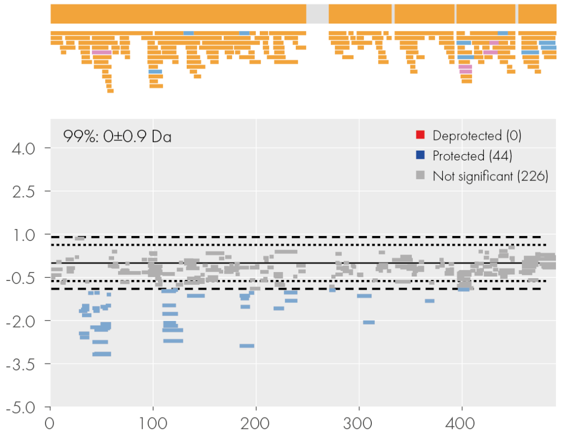


### 21. E206Q&D27N - D27N+Xylose

Common: 98.98% / XylE D27N_Xylose: 98.98% / XylE E206Q_D27N: 98.98%

Common: 230 XylE D27N_Xylose: 1 XylE E206Q_D27N: 0

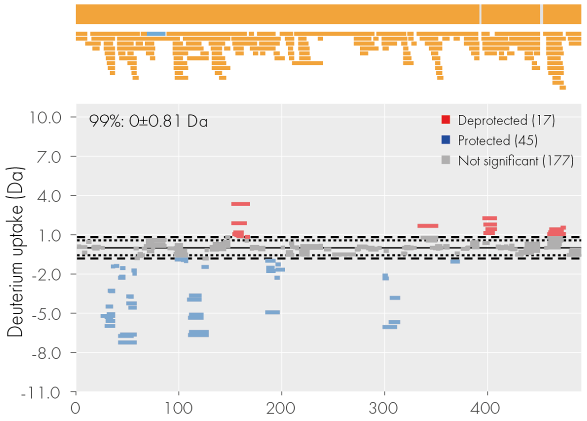


### 22. E206Q&D27N+Xylose - D27N+Xylose

Common: 97.76% / XylE D27N+Xylose: 97.76% / E206Q&D27N+Xylose: 97.76%

Common: 204 XylE D27N+Xylose: 1 E206Q&D27N+Xylose: 1

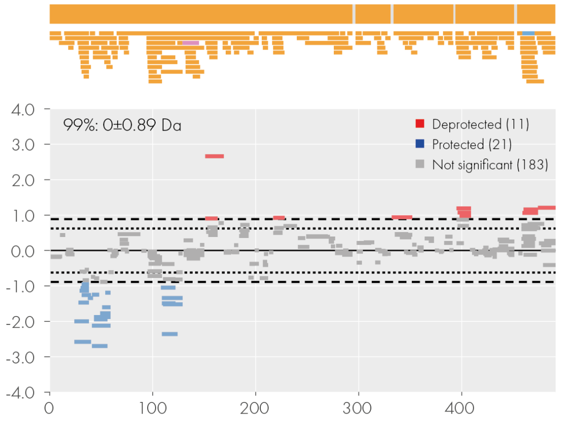


### 23. E206Q+Xylose - E206Q

Common: 97.35% / XylE E206Q: 97.35% / XylE E206Q+Xylose: 97.96%

Common: 219 XylE E206Q: 1 XylE E206Q+Xylose: 3

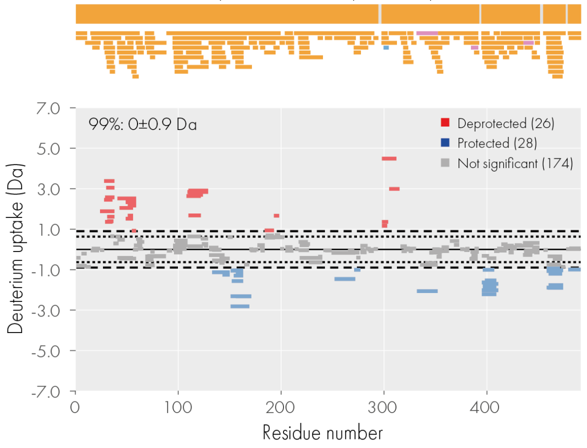


## 24. E206Q&D27N - E206Q

Common: 97.35% / XylE E206Q: 97.35% / XylE E206Q&D27N: 97.35%

Common: 220 XylE E206Q: 0 XylE E206Q&D27N: 1

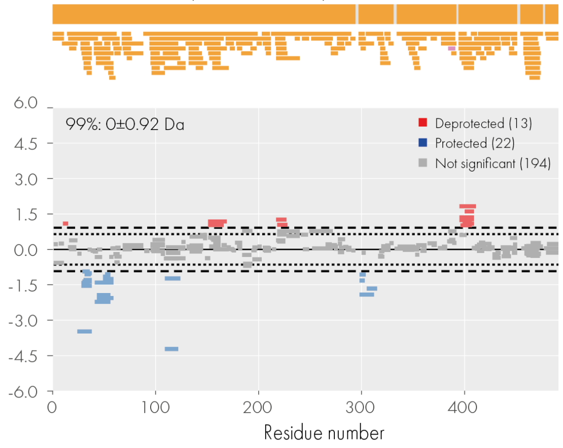

**25. E206Q&D27N+Xylose - E206Q**

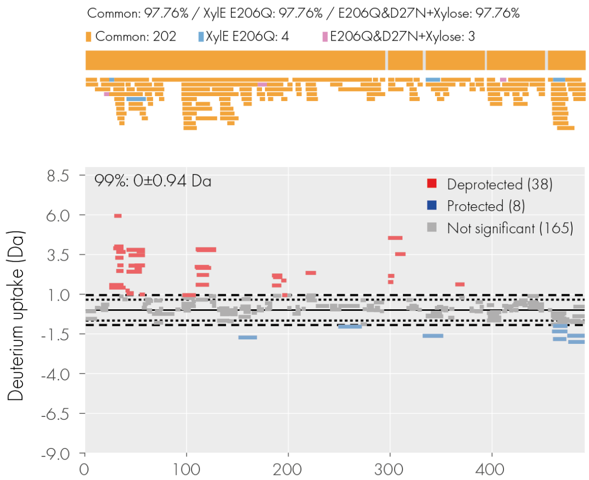


**26. E206Q+Xylose - E206Q&D27N**

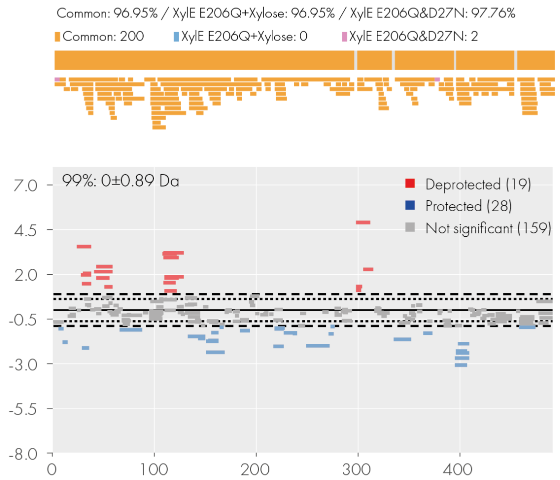


**27. E206Q&D27N +Xylose - E206Q+Xylose**

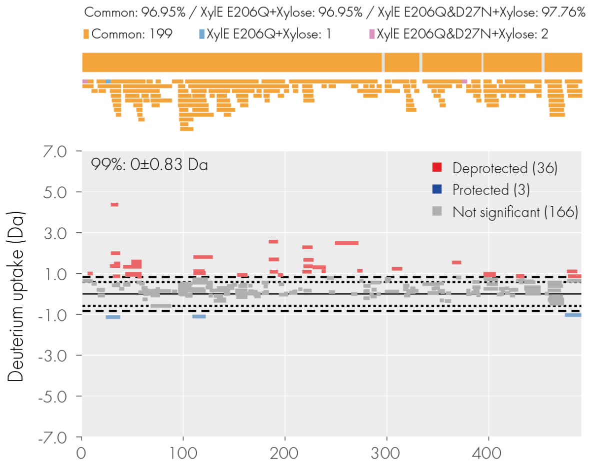


**28. E206Q&D27N +Xylose - E206Q&D27N**

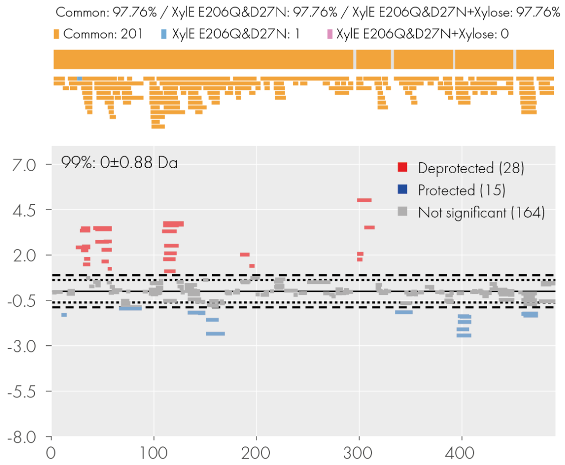


**29. WT+Glucose - WT**

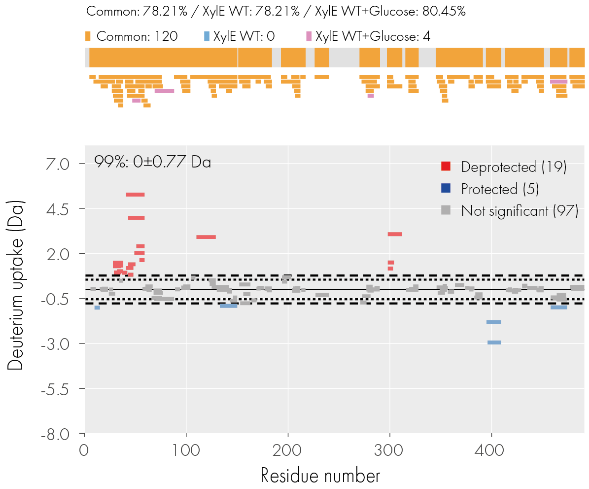


**30. D27N+Glucose - WT**

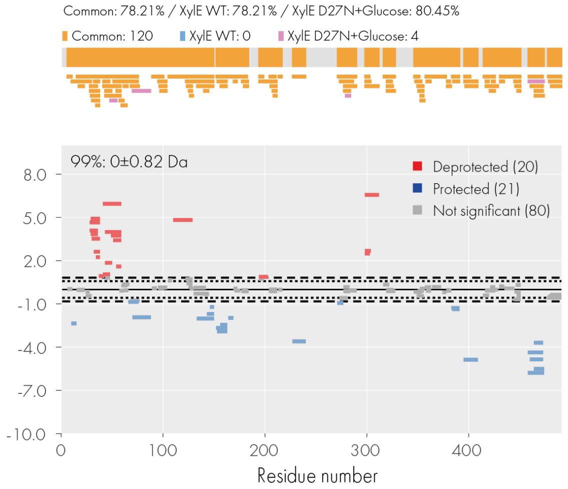

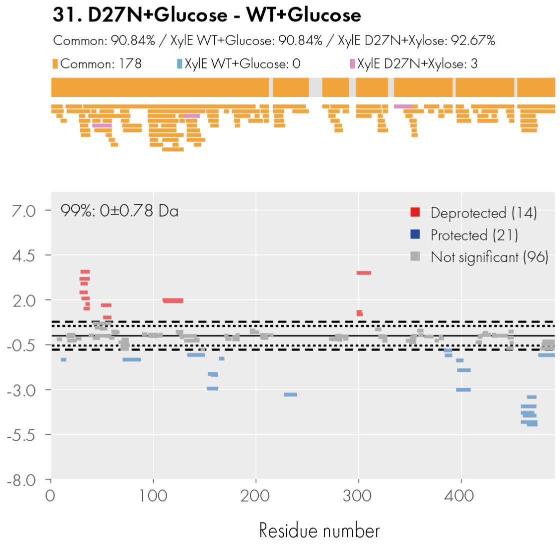


**Figure S4. Woods plots and comparative sequence coverage maps obtained from differential HDX of XylE.** Each bar represents a single peptide with peptide length indicated by the bar length. Common peptide between two different protein states are indicated as orange, unique peptides are in blue and pink separately.

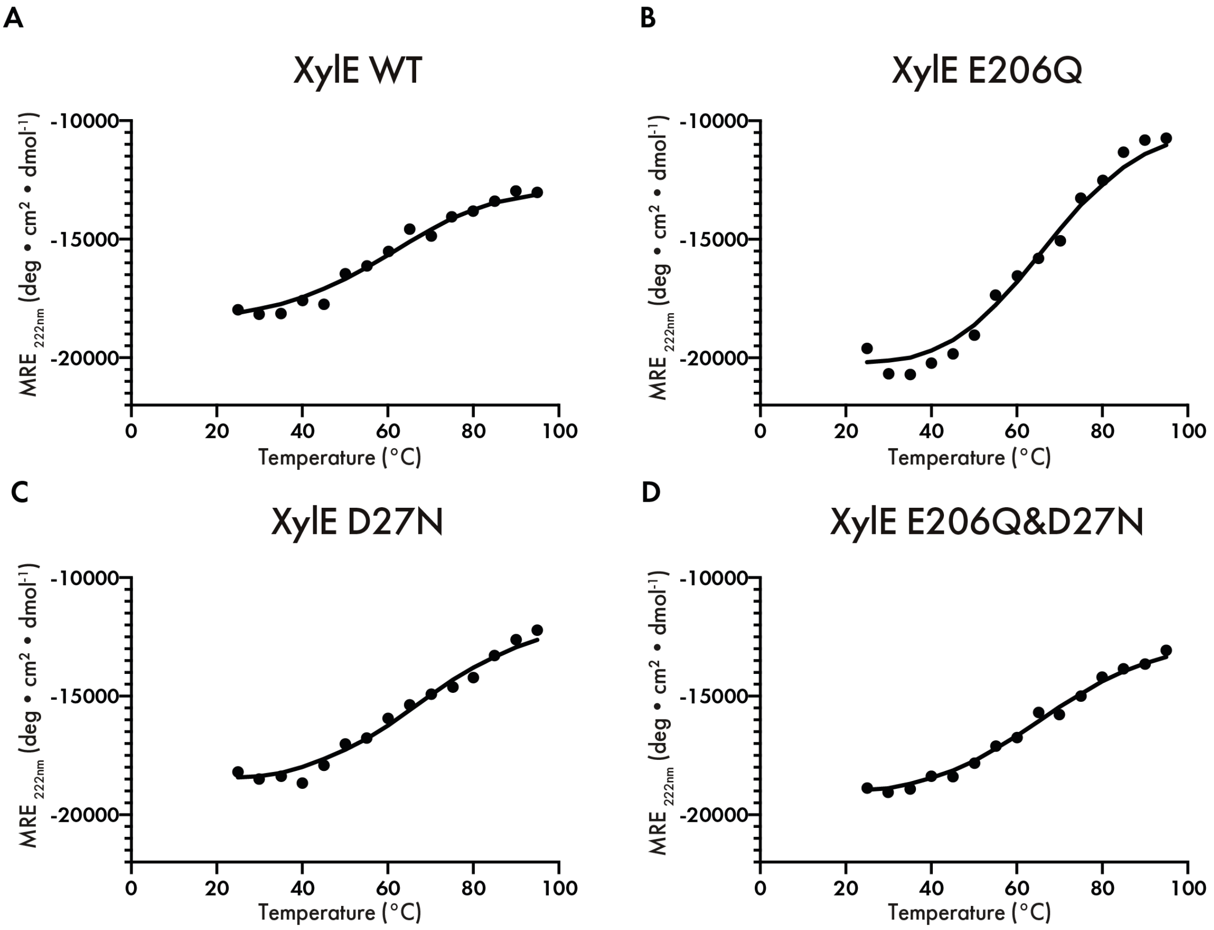


**Figure S5. Thermal denaturation Circular Dichroism of XylE wild type, E206Q, D27N and E206Q&D27N**. (**A-D).** Measurements of XylE WT, E206Q, D27N, and E206Q&D27N on the changes in MRE (Mean Residue Ellipticity) at fixed wavelength 222 nm from 25 ℃ to 95 ℃.

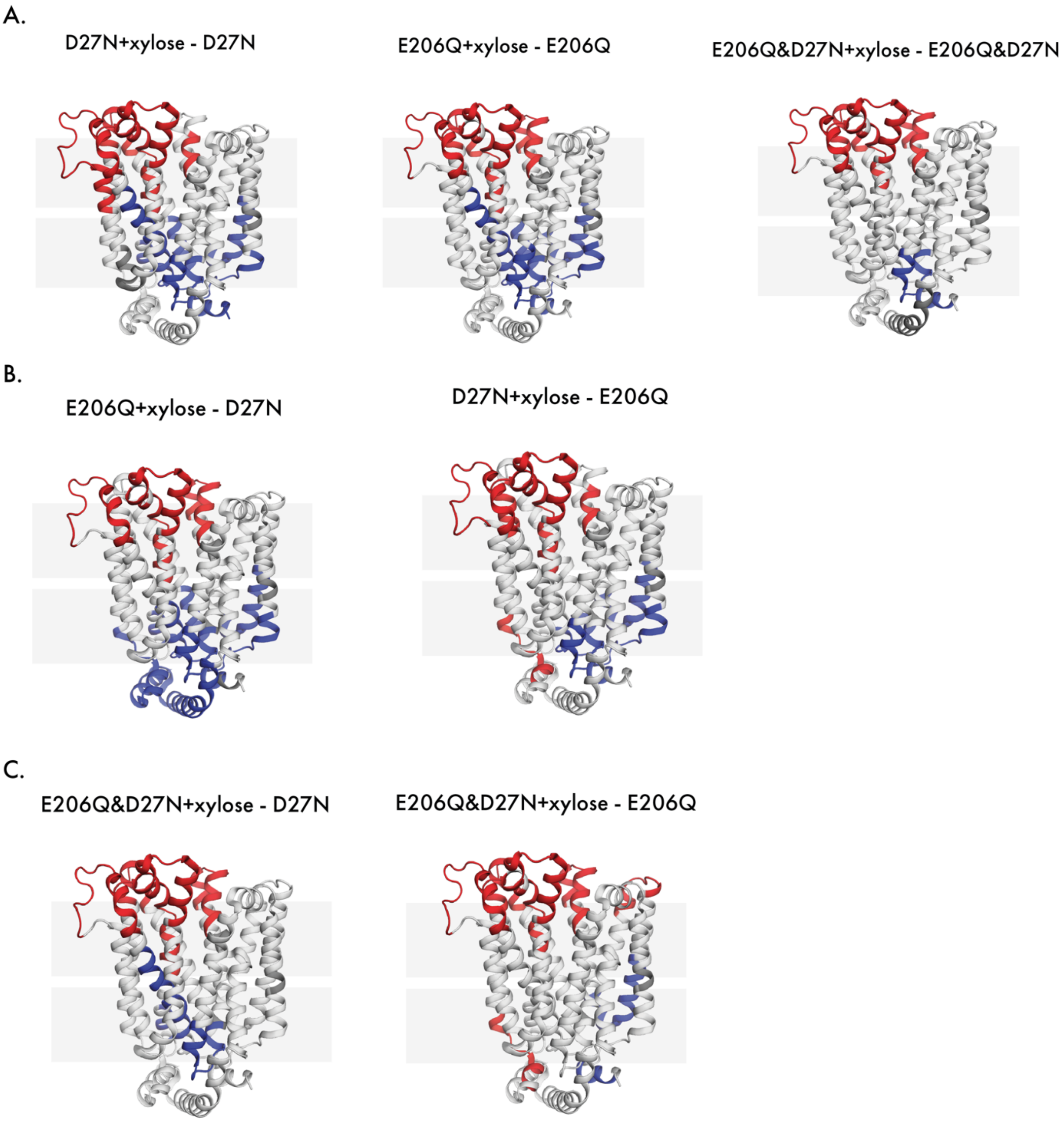


**Figure S6. Conformational equilibrium towards outward-facing through substrate binding. (A).** Differential deuterium uptake pattern of binding of the substrate to a singly protonated state minus the same singly protonated state without substrate. (**B).** Differential deuterium uptake pattern of binding of the substrate to a singly protonated state minus a different singly protonated state. (**C).** Differential deuterium uptake pattern of binding of the substrate to a doubly protonated state minus a singly protonated state. Figures are plotted onto the 3D protein structure (PDB: 4GBY) by PyMol. Blue and red regions suggest a relatively negative (protected) and a positive (deprotected) deuterium uptake pattern respectively.

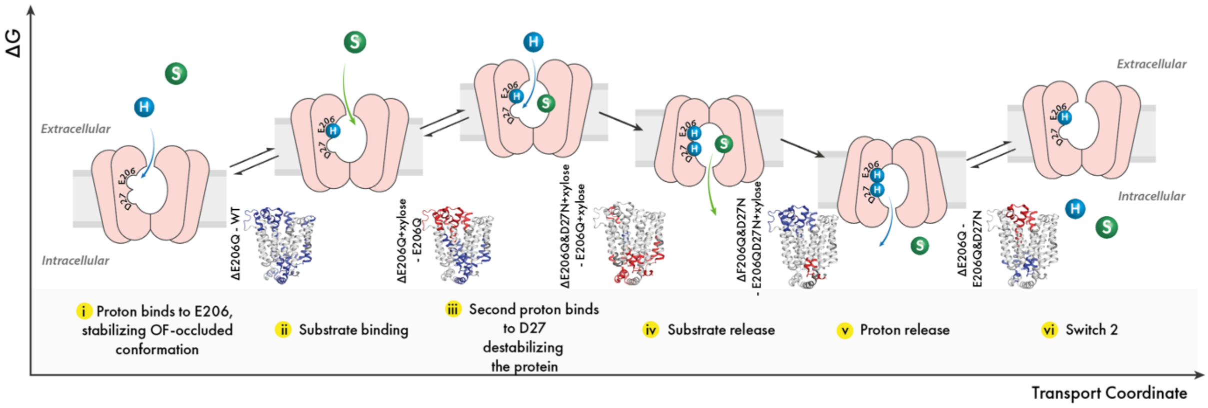


**Figure S7. Proposed model of energy landscape of XylE active transport. (i)** E206 is always protonated. **(ii)** Comparison of d(E206Q+xylose – E206Q) indicates binding of substrates leads to OF conformation. **(iii)** Comparison of d(E206Q&D27N+xylose – E206Q+xylose) indicates that another proton binds to D27 triggers the conformational switch. **(iv)** Comparison of d(E206Q&D27N - E206Q&D27N+xylose) indicates that substrate release shifts conformational equilibrium towards inward-facing. **(v)** and **(vi)** Comparison of d(E206Q – E206Q&D27N) indicates once proton released, it goes to next round of switch.
